# ALS mice carrying pathological mutant TDP-43, but not mutant FUS, display axonal transport defects *in vivo*

**DOI:** 10.1101/438812

**Authors:** James N. Sleigh, Andrew P. Tosolini, David Gordon, Anny Devoy, Pietro Fratta, Elizabeth M. C. Fisher, Kevin Talbot, Giampietro Schiavo

## Abstract

Amyotrophic lateral sclerosis (ALS) is a fatal, progressive neurodegenerative disease resulting from a complex interplay between genetics and environment. Impairments in the basic neuronal process of axonal transport have been identified in several ALS models. However, *in vivo* evidence of early/pre-symptomatic deficiencies in neuronal cargo trafficking remains limited, thus the pathogenic importance of axonal transport to the ALS disease spectrum remains to be fully resolved. We therefore analysed the *in vivo* dynamics of retrogradely transported, neurotrophin-containing signalling endosomes in motor neuron axons of two new mouse models of ALS that have mutations in different RNA processing genes (*Tardbp* and *Fus*). TDP-43^M337V^ mice, which show neuromuscular pathology but no overt motor neuron loss, displayed *in vivo* perturbations in axonal transport that manifested between 1.5 and 3 months and preceded motor symptom onset. In contrast, signalling endosome transport remained largely unaffected in mutant Fus^Δ14/+^ mice, despite 20% motor neuron loss. These findings indicate that deficiencies in retrograde neurotrophin signalling and axonal transport are not common to all ALS-linked genes, and that there are inherent and mechanistic distinctions in the pathogenesis of ALS caused by mutations in different RNA processing genes.

## Introduction

Amyotrophic lateral sclerosis (ALS) is a rapidly progressive neurodegenerative disorder that results from upper and lower motor neuron loss, leading to muscle wasting, atrophy, and, ultimately, death most often due to respiratory failure (Brown and Al-Chalabi 2017). Treatment options for ALS patients are severely limited, but gene therapy approaches hold great promise (Tosolini and Sleigh 2017). ALS is thought to manifest through a multi-step process encompassing additive effects from genetic predispositions and environmental insults (Al-Chalabi and Hardiman 2013); however, ≈10% of cases show clear monogenic heritability (familial ALS, fALS), while known causative genetic mutations underlie ≈68% of fALS and ≈11% of the remaining sporadic cases of ALS (Renton *et al.* 2014). Mutations in numerous genes are reproducibly linked to the disease, the four most common of which, in ascending order, are dominant mutations in *fused in sarcoma* (*FUS*), *transactive-region DNA binding protein* (*TARDBP* encoding TDP-43) and *superoxide dismutase 1* (*SOD1*), and large, intronic hexanucleotide repeat expansions in *chromosome 9 open reading frame 72* (*C9orf72*) (Brown and Al-Chalabi 2017).

Many genes associated with ALS encode proteins important in all cells, and as such, it remains unknown why motor neurons and certain brain regions, such as the frontotemporal cortex, are selectively affected. Nonetheless, impairments in cytoskeletal dynamics and axonal transport are emerging as a central theme based on ALS-linked gene function (Clark *et al.* 2016; De Vos & Hafezparast 2017). Axonal transport is the essential, bi-directional process whereby cargoes (e.g. organelles and proteins) are actively transported from one end of an axon to the other along microtubules (Maday *et al.* 2014). Anterograde transport, which is from the cell body to axon tip, is dependent on the kinesin family of molecular motors, while cytoplasmic dynein is responsible for retrograde axonal transport in the opposite direction. Patient post-mortem studies provided the first evidence for involvement of impaired transport in ALS, which has since been consolidated by results from a plethora of disease models implicating various cargoes (De Vos & Hafezparast 2017). Transport deficits have been linked to all four major ALS genes through *in vitro*, *ex vivo* and *Drosophila melanogaster* experiments; however, these models do not necessarily replicate the complex environment found in mammals required for efficient, rapid axonal transport (Sleigh *et al.* 2017). *In vivo* results from mammals in which individual cargoes are tracked in real time, rather than *en masse*, have been generated in SOD1^G93A^ and TDP-43^A315T^ ALS mice (**Supplementary Table 1**) (Bilsland *et al.* 2010; Magrané *et al.* 2014; Gibbs *et al.* 2018). Axonal transport is disrupted in both models at early disease stages, consistent with a potential causative role in neuromuscular dysfunction and motor neuron degeneration; nonetheless, it is unclear whether these ALS mice, which express disease-causing mutant proteins at supra-physiological levels, are reflective of the full disease spectrum.

We have thus performed pseudolongitudinal assessments of *in vivo* axonal transport in two recently engineered mouse models of ALS with mutations in genes encoding DNA/RNA-binding proteins instrumental to RNA processing, TDP-43 (Gordon *et al.* 2018) and Fus (Devoy *et al.* 2018). Transgenic TDP-43^M337V^ and humanised, knock-in Fus^Δ14/+^ mice, which both express mutant protein at physiologically-relevant levels, have been used to assess signalling endosome trafficking along peripheral nerve axons to address the importance of altered axonal transport to ALS neuropathology.

## Materials and Methods

### Animals

All mouse handling and experiments were performed under license from the United Kingdom Home Office in accordance with the Animals (Scientific Procedures) Act (1986), and approved by the University College London – Institute of Neurology Ethics Committee. Tg(Chat-EGFP)GH293Gsat/Mmucd mice (MMRRC Stock Number 000296-UCD), referred to as ChAT-eGFP mice, were maintained and imaged as heterozygotes on a CD-1 background. B6;129S6-Gt(ROSA)26Sor^m1(TARDBP*M337V/Ypet)Tlbt^/J (WT and M337V TDP43, Jackson Laboratory strain #029266, https://www.jax.org/strain/029266) and B6N;B6J-Fustm1Emcf/H Fus^+/+^ and Fus^Δ14/+^) mice were maintained on a C57BL/6 background and genotyped as detailed previously (Devoy *et al.* 2018; Gordon *et al.* 2018). ChAT-eGFP mice used for motor versus sensory analyses were 79-134 days old. Non-transgenic (NTg) control and TDP43 mice sacrificed for 1.5, 3, and 9 month time points were 55-57, 102-125, and 249-71 days old, respectively. Fus mice sacrificed for 3, 12, and 18 month time points were 104-115, 365-368, and 568-588 days old, respectively.

### Axonal transport analysis

*In vivo* kinetics of signaling endosomes labelled with atoxic binding fragment of tetanus neurotoxin (H_C_T) were assessed as previously described (Gibbs *et al.* 2016, Sleigh and Schiavo 2016). Briefly, H_C_T^441^ (residues 875-1315) fluorescently labelled with AlexaFluor555 C_2_ maleimide (Life Technologies, A-20346) was injected into the motor end plate region of gastrocnemius and tibialis anterior muscles of the right leg as per Mohan *et al.* (2014). 5 μg of H_C_T, pre-mixed with 25 ng recombinant human BDNF (Peprotech, 450-02), was injected per muscle in a volume of ≈1.5 μl. 4-8 h post-injection, the right sciatic nerve was exposed in isofluorane-anaesthetised mice and imaged on an inverted LSM780 laser scanning microscope (Zeiss) within an environmental chamber at 37°C. Endosomes were imaged in at least three distinct axons per animal, and images acquired every 2.4-3.2 s. Image series were converted into .avi files and individual endosome dynamics manually tracked using Kinetic Imaging Software. Endosomes were included in the analysis if they could be observed for ≥5 consecutive frames, and did not pause for >10 consecutive images. An endosome was considered to have paused if it remained in the same position for two consecutive images. All individual frame-to-frame step speeds are included in the presented speed frequency histograms (485.8 α 13.1 frame-to-frame speeds per animal were calculated). To determine the mean endosome speed per animal, the speeds of individual endosomes were calculated and averaged (51.6 α 0.7 endosomes per animal were tracked). The fastest endosome speed per animal is reported as the maximum speed. At least ten endosomes from at least three individual, thick axons were assessed per animal within 1 h of initiating terminal anaesthesia.

### Axon calibre analysis

Axon calibres were determined from images taken for endosome transport analyses by measuring the distance between the upper and lower margins of transported fluorescent signalling endosomes at 90° from the direction of transport. A minimum of ten measurements were made along the length of the axon to calculate average widths per axon, and three different axons per animal were used to calculate a per animal mean width.

### Statistical analysis

Data were assumed to be normally distributed unless evidence to the contrary could be provided by the D’Agostino and Pearson omnibus normality test. Normally distributed data were statistically analysed using a *t*-test or one-way analysis of variance (ANOVA) with Dunnett’s multiple comparisons test, and non-normally distributed data with a Mann-Whitney *U* test or Kruskal-Wallis test with Dunn’s multiple comparisons test. Paired *t*-tests were used to compare transport kinetics in ChAT^+^ versus ChAT^−^ axons as data were generated from the same animals. Endosomes were tracked from videos in which the genotype of the animal was blinded. All tests were two-tailed and an α-level of *P* < 0.05 was used to determine significance. GraphPad Prism 6 software was used for all statistical analyses and figure production. Means ± standard error of the mean are plotted for all graphs and are the statistics reported in the main text. No significant differences in transport were observed between sexes for any genotype (**Supplementary Figure 4E-H**, and data not shown), so data from males and females were pooled in all presented analyses.

**Figure 1.**
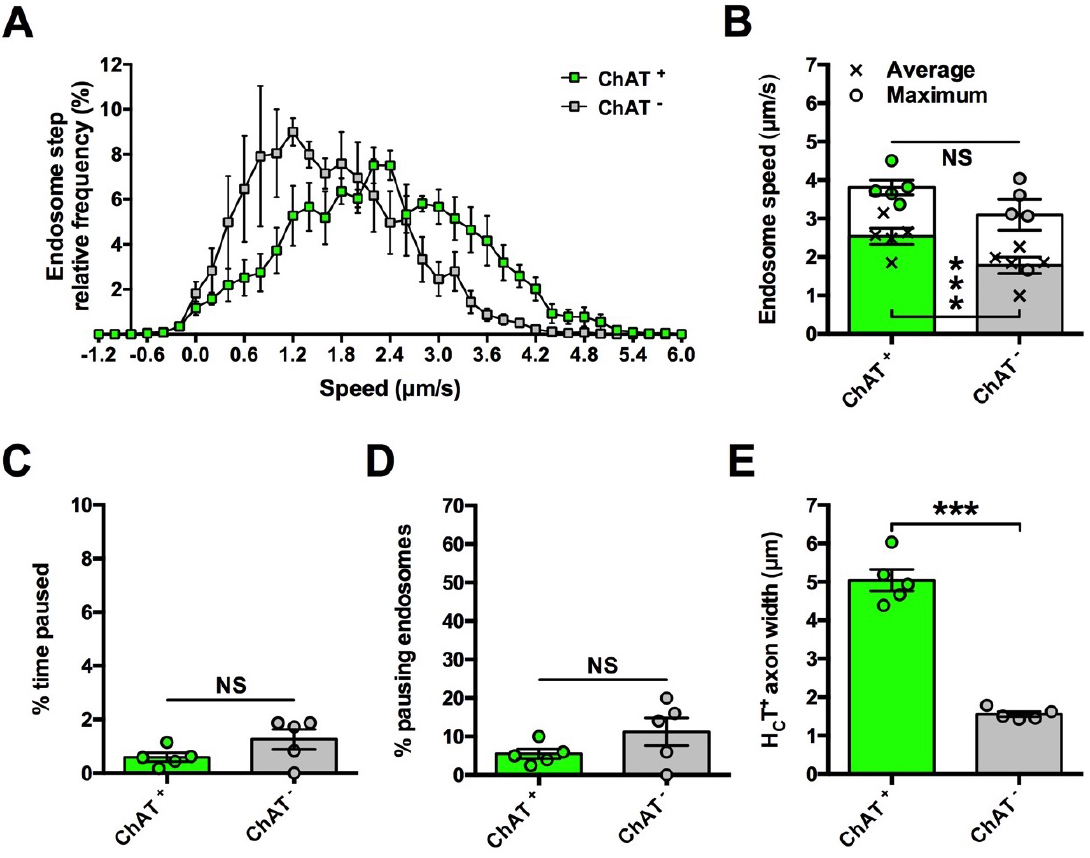
Retrograde axonal transport of signalling endosomes is faster in motor neurons than sensory neurons. (**A**) Speed distribution curves of signalling endosome frame-to-frame movements in motor (ChAT^+^, green) and sensory axons (ChAT^−^, grey) indicate that axonal transport is faster in motor neurons. (**B**) Mean (crosses), but not maximum (circles), endosome transport is faster in motor neurons when calculated per animal. (**C** and **D**) There is no difference between motor and sensory nerves in the percentage of time endosomes paused for (C) or the percentage of endosomes that paused (D). (**E**) H_C_T-containing axons that are ChAT^+^ have a larger calibre than ChAT^−^ axons. *** *P* < 0.001; NS, not significant, paired *t*-test. *n* = 5. See also **Supplementary Fig. 1**.

**Figure 2.**
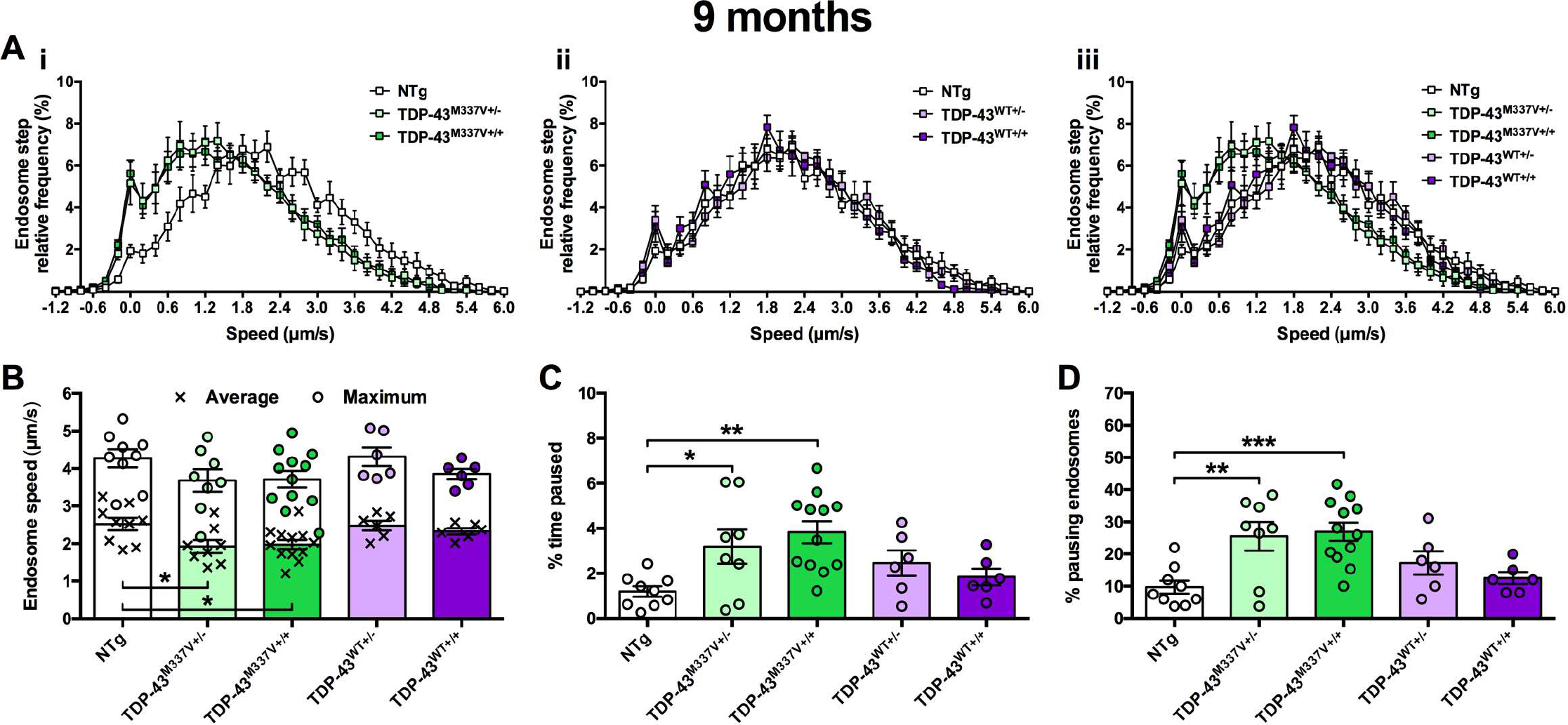
Mutant TDP mice display perturbed axonal transport of endosomes at 9 months. (**A**) Endosome frame-to-frame speed distribution curves indicate that heterozygous and homozygous TDP-43^M337V^ mice (i, iii, green) transport endosomes more slowly than control, non-transgenic (NTg) mice (white), whereas transport is unaffected in TDP-43^WT^ controls (ii, iii, purple). (**B**) Mean endosomal speeds (crosses) are significantly reduced in TDP-43^M337V+/-^ and TDP-43^M337V+/+^ mice, but not TDP-43^WT^ controls (*P* = 0.011, one-way ANOVA), while maximum speeds (circles) remained unchanged (*P* = 0.223, one-way ANOVA). (**C**) Endosomes in mutant TDP-43^M337V^ heterozygous and homozygous mice paused for longer periods of time compared to NTg mice (*P* = 0.004, one-way ANOVA). (**D**) TDP-43^M337V^ mice had a greater percentage of endosomes that paused (*P* < 0.001, one-way ANOVA). * *P* < 0.05; ** *P* < 0.01; *** *P* < 0.001, Dunnett’s multiple comparisons test. *n* = 6-12. See also **Supplementary Fig. 2** and **4**.

## Results

### Imaging *in vivo* axonal transport in motor neurons

To assess *in vivo* dynamics of axonal transport, we used a fluorescently-labelled binding fragment of tetanus neurotoxin (H_C_T), which is retrogradely transported along axons within neurotrophin-containing signalling endosomes towards neuronal cell bodies (Surana *et al.* 2018; Villarroel-Campos *et al*. 2018). Impairments in this long-range neurotrophic signalling have been implicated in several neurodegenerative conditions including ALS (Bronfman *et al.* 2007). By injecting H_C_T into the gastrocnemius and tibialis anterior muscles of the lower leg, and exposing the sciatic nerve at thigh-level 4-8 h post-injection, individual, fluorescently-labelled endosomes being retrogradely transported can be imaged and tracked in the peripheral nerve axons of live, anaesthetised mice (Gibbs *et al.* 2016).

Post-intramuscular injection, ≈80% of H_C_T^+^ axons stain for choline acetyltransferase (ChAT) (Bilsland *et al.* 2010), suggesting that the probe is preferentially transported in motor neurons. Nevertheless, assessing transport in a mixed motor and sensory population may weaken the ability to identify motor-specific trafficking perturbations. Therefore, before analysing transport in ALS mice, we compared endosome dynamics in motor versus sensory neurons using ChAT.eGFP mice, which permit visual differentiation of peripheral nerve types because motor axons are specifically labelled with eGFP. Mean endosome transport speeds were greater in ChAT^+^ motor neurons compared to ChAT^−^ sensory neurons (**Fig. 1A** and **B**), and this was not due to pausing differences (**Fig. 1C** and **D**). Moreover, motor axons had clearly larger calibres than sensory axons (**Fig. 1E**). This suggests that by imaging thicker axons, H_C_T transport can be measured in motor neurons with greater certainty than if randomly selecting an axon (*i.e.* >80%, Bilsland *et al.* 2010). To confirm this, transport dynamics were compared between ChAT^+^ axons and thicker axons from non-fluorescent control mice, and no differences were observed (**Supplementary Fig. 1**). The bell-shaped, rather than bi-modal, speed frequency distribution generated from non-fluorescent mice (**Supplementary Fig. 1A**) indicates that *in vivo* axonal transport of endosomes can be assessed predominantly in motor neurons by selecting large calibre axons. This approach was thus used to analyse axonal transport in ALS mice.

### *In vivo* axonal transport is pre-symptomatically impaired in mutant TDP43 mice

Recently reported transgenic TDP-43^M337V^ mice display an impairment in motor function and neuromuscular junction abnormalities beginning at 9 months in homozygous mutants without motor neuron loss up to 12 months (Gordon *et al.* 2018). We therefore first assessed retrograde transport of signalling endosomes at 9 months of age in heterozygous and homozygous TDP-43^M337V^ and TDP-43^WT^ mice and non-transgenic (NTg) controls (**Fig. 2**). The frequency histograms of frame-to-frame endosome speeds of both TDP-43^M337V+/−^ and TDP-43^M337V+/+^ animals are shifted to the left compared to NTg mice, indicative of slower transport, whereas TDP-43^WT^ transport was unaffected since it perfectly overlaps with the curve obtained using NTg controls (**Fig. 2A**). When statistically compared, both mutants showed a significant reduction in mean endosome speed (**Fig. 2B**), which was at least partially due to increased pausing (**Fig. 2C** and **D**). Mutant TDP-43 mice do not show clear behavioural phenotypes at 3 months (Gordon *et al*. 2018); we therefore assessed transport at this early time point to see whether axonal transport defects precede symptom onset and thus may contribute to motor neuron pathology. Indeed, a similar deficiency in mutant TDP-43 transport was observed at 3 months, while TDP-43^WT^ transport remained unperturbed (**Fig. 3A-D**). Finally, to determine at what stage transport becomes affected, we assessed endosomal trafficking at 1.5 months in TDP-43^M337V+/+^ and TDP-43^WT+/+^ mice. We found no difference between genotypes (**Fig. 3E-H**) or from NTg control mice (not shown). These data indicate that TDP-43^M337V^, but not TDP-43^WT^, mice display a pre-symptomatic *in vivo* impairment in axonal transport of signalling endosomes that manifests between 1.5 and 3 months of age (**Supplementary Fig. 2**).

**Figure 3.**
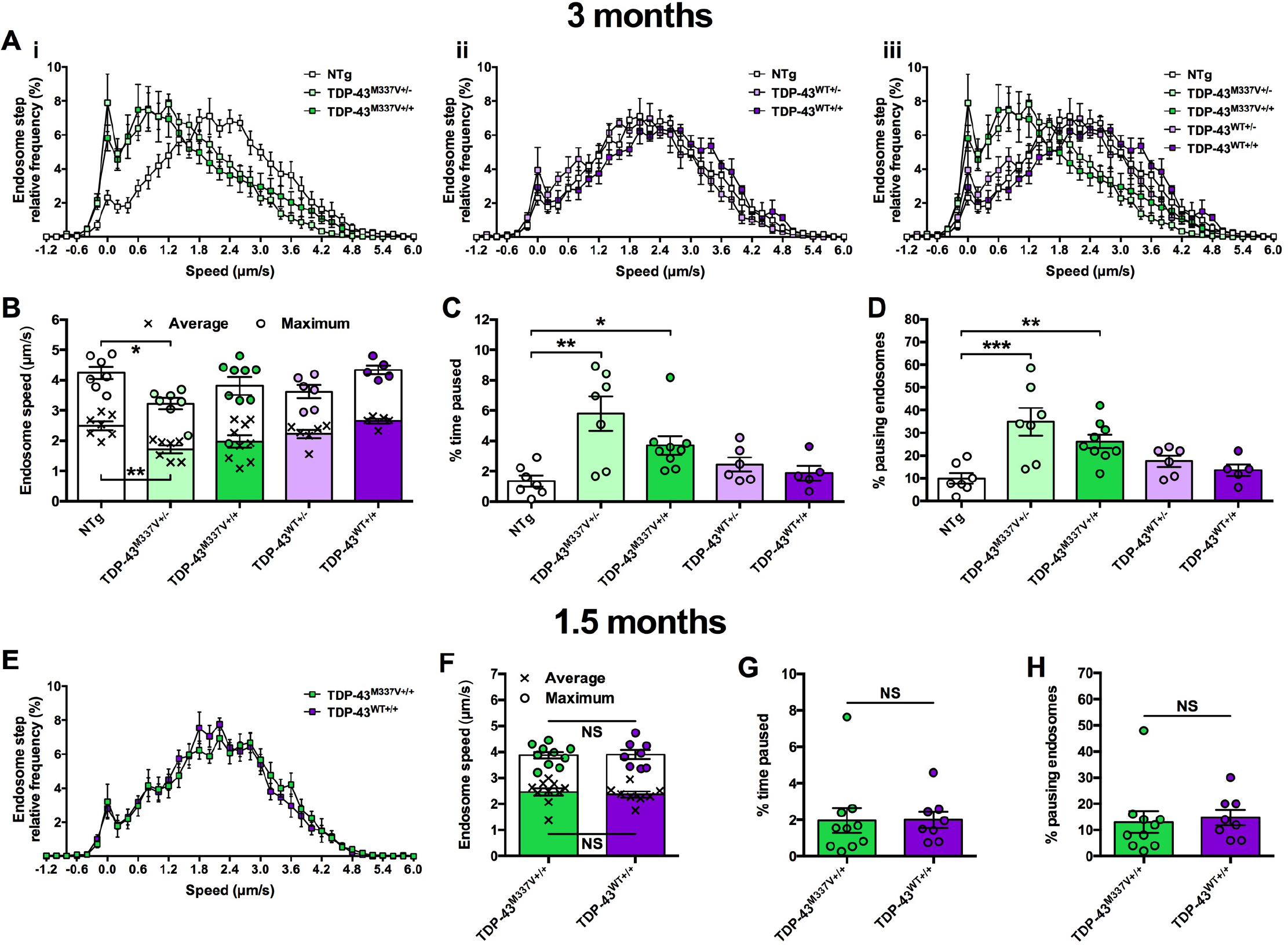
TDP-43^M337V^ transport disruption occurs between 1.5 and 3 months. (**A**) Endosome frame-to-frame speed distribution curves show that at 3 months TDP-43^M337V^ (i, iii, green), but not TDP-43^WT^ (ii, iii, purple), mice transport endosomes more slowly than control, non-transgenic (NTg) mice (white). (**B**) Mean (crosses, *P* = 0.003, one-way ANOVA) and maximum (circles, *P* = 0.022, one-way ANOVA) endosomal speeds are significantly reduced in TDP-43^M337V+/-^ mice. (**C** and **D**) TDP-43^M337V^ heterozygotes and homozygotes show increased endosome pausing as assessed by calculating the percentage of time paused (C, *P* = 0.004, Kruskal-Wallis test) and the percentage of pausing endosomes (D, *P* < 0.001, one-way ANOVA). (**E-H**) At 1.5 months there is no difference in endosome transport speeds or pausing between TDP-43^M337V+/+^ (green) and TDP-43^WT+/+^ (purple) mice, suggesting that transport disruption occurs between 1.5 and 3 months. Presented 1.5 month data are not significantly different from NTg control (not shown). * *P* < 0.05; ** *P* < 0.01; *** *P* < 0.001, Dunnett’s/Dunn’s multiple comparisons test. NS, not significant, unpaired *t*-test/Mann-Whitney *U* test. *n* = 5-10. See also **Supplementary Fig. 2** and **4**.

### Axonal transport remains largely unaffected in mutant Fus mice even at late stages

Deficient *in vivo* signalling endosome trafficking has now been observed in SOD1^G93A^ mice (Bilsland *et al.* 2010; Gibbs *et al.* 2018) and the TDP-43^M337V^ model reported here. To assess whether this phenotype is common to mouse models of ALS, we assessed *in vivo* transport in knock-in mutant Fus^Δ14/+^ mice. This model displays loss of neuromuscular integrity and progressive degeneration of lumbar spinal motor neurons; at 3 months, mutant Fus mice show no motor neuron loss, which becomes overt by 12 (14% reduction) and 18 (20% reduction) months of age (Devoy *et al.* 2018). We therefore assessed endosome transport at 3, 12, and 18 months in this novel ALS model (**Fig. 4**). At 3 and 12 months, there was no significant difference in endosome kinetics (**Fig. 4A-H**), and, despite minor increases in pausing (**Fig. 4K** and **L**), there was no significant change in signalling endosome mean or maximum speeds at the late disease stage of 18 months (**Fig. 4I** and **J**). Consistent with this, no significant changes in transport were observed across time points for Fus^+/+^ or Fus^Δ14/+^ mice, although Fus mutants perhaps show a subtle progressive decline as a secondary consequence of neurodegeneration (**Supplementary Fig. 3**).

**Figure 4.**
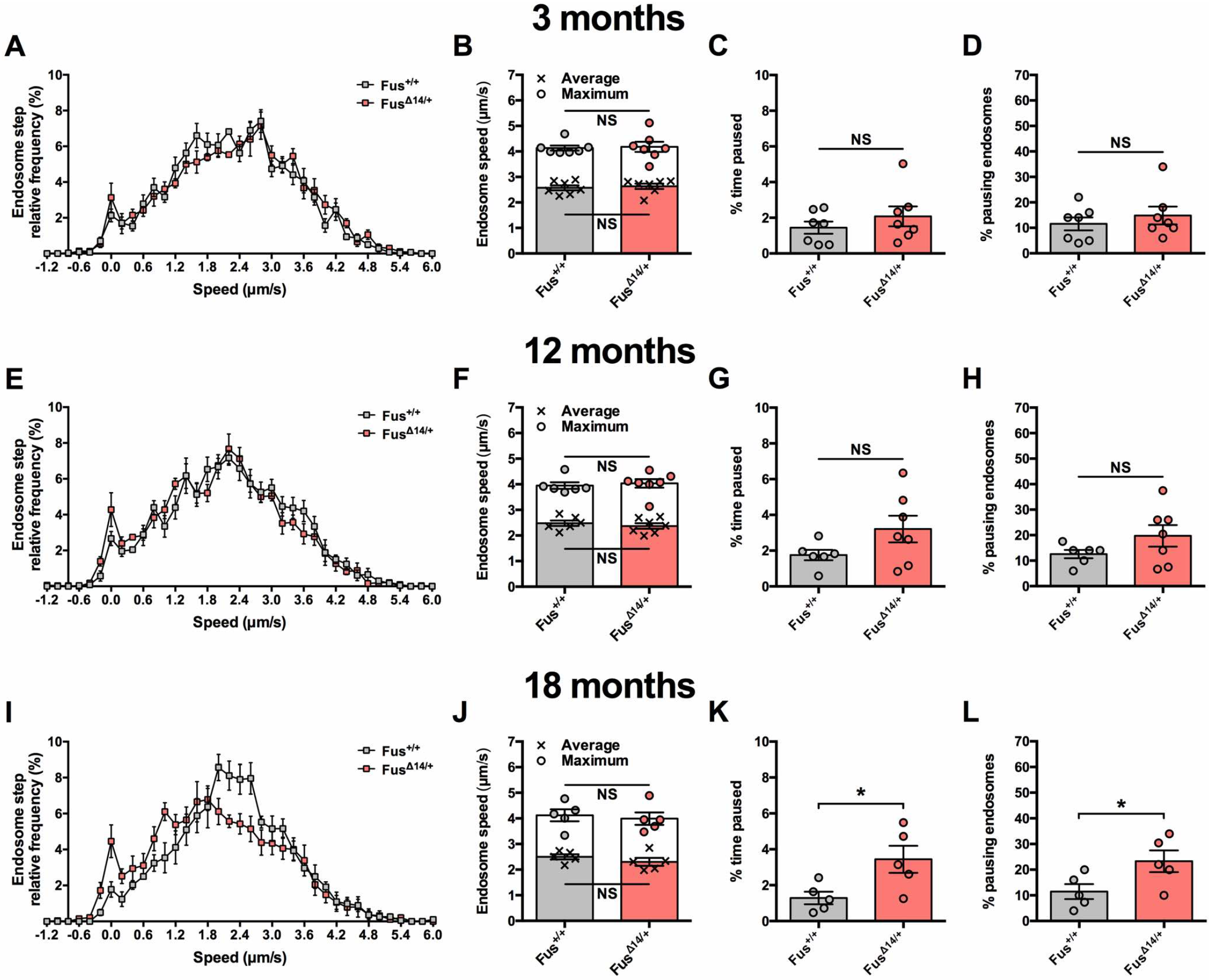
Fus^Δ14/+^ mice display a minor impairment in axonal transport of endosomes, but only at a late disease stage. (**A-H**) The axonal dynamics of signalling endosomes are similar between Fus^+/+^ (grey) and Fus^Δ14/+^ (red) mice at 3 months (A-D) and 12 months (E-H) of age. (**I-L**) At 18 months, retrograde axonal transport speed of endosomes is unaffected in Fus^Δ14/+^ mice (I and J); however, mutant Fus mice do show a mild increase in the percentage of time that endosomes pause for (K) and the percentage of endosomes that paused (L). * *P* < 0.05; NS, not significant, unpaired *t*-test/Mann-Whitney *U* test. *n* = 5-7. See also **Supplementary Fig. 3** and **4**.

We have previously shown that endosome transport remains stable in wild-type mice from 1 to 13-14 months of age (Sleigh & Schiavo 2016), suggesting that a natural, aging-related decline in transport does not compound the mutant TDP-43 transport defect. To ensure that this remains true up to 18 months, we compared axonal transport in all control mice aged 3-18 months. There were no significant changes in cargo dynamics (**Supplementary Fig. 4**), suggesting that the mild pausing defect of 18 month-old mutant Fus mice is unlikely to be a direct consequence of aging, rather a secondary effect of a degenerating motor system, and that axonal transport of signalling endosomes remains unaltered in wild-type mice up to 18 months.

## Discussion

For the first time, we show that an ALS mouse model of mutant TDP-43 displays a pre-symptomatic, *in vivo* deficit in axonal transport of signalling endosomes in motor axons, which may contribute to motor function deficits and impaired neuromuscular integrity. This defect is specific to the M337V mutation, as TDP-43^WT^ protein, which is expressed at a similar low level as TDP-43^M337V^ relative to endogenous mouse TDP-43 (Gordon *et al.* 2018), had no effect on transport. This adds to the impaired mitochondrial transport reported in TDP-43^A315T^ mice and defective mitochondria and signalling endosome trafficking in SOD1^G93A^ mice (**Supplementary Table 1**). ALS-linked mutations in *SOD1* and *TARDBP* may thus cause early/pre-symptomatic, generalised defects in axonal transport in motor neurons (rather than cargo-specific deficits), leading to dysfunction and degeneration (Gordon *et al.* 2018). This may be caused by non-selective impairments in the cytoskeleton or molecular motor proteins in motor neurons, or perhaps by aberrant binding of mutant ALS proteins to multiple motor protein complexes (Zhang *et al.* 2007; Tateno *et al.* 2009); however, this will have to be directly confirmed in the TDP-43^M337V^ model.

Contrastingly, Fus^Δ14/+^ endosome transport remained largely unaffected even during latter disease stages, despite a 20% loss of spinal cord motor neurons (Devoy *et al.* 2018), confirming that degenerating axons do not always have altered transport kinetics (Malik *et al.* 2011). This implies that transport disturbances are not necessarily a non-specific bi-product of neurodegeneration, at least during earlier disease stages, and thus emphasises the specificity of transport defects in mutant TDP-43 and SOD1 mice. However, it remains possible that rapid waves of degeneration of motor neuron subtypes occur in Fus^Δ14/+^ mice such that any preceding defect in transport was missed (Nijssen *et al.* 2017). Alternatively, mutant Fus mice may display cargo-specific (e.g. mitochondria) or anterograde transport defects, which occur in other ALS models (Alami *et al.* 2014; Baldwin *et al.* 2016), thus additional cargoes should also be assessed in Fus^Δ14/+^ mice. Nevertheless, together our findings indicate that pre-symptomatic deficiencies in retrograde axonal transport of neurotrophin-containing signalling endosomes may not be common to all ALS-linked genes, and that there are inherent distinctions in the pathogenesis of ALS caused by mutations in different RNA processing genes. While TDP-43 and FUS are both RNA/DNA-binding proteins that process RNA predominantly in the nucleus, they regulate the expression/splicing of largely distinct gene sets (Colombrita *et al.* 2012; Lagier-Tourenne *et al.* 2012) and show neuropathological idiosyncrasies when mutated (Bäumer *et al.* 2010), which could account for the transport discrepancy.

Disruptions in axonal transport have been linked to the M337V TARDBP mutation in a range of *in vitro* and *Drosophila* larval models (Wang *et al*. 2013; Alami *et al.* 2014; Baldwin *et al*. 2016), and, while the severe, frameshift FUS mutation modelled in Fus^Δ14/+^ mice has not previously been assessed, transport perturbations have been reported in several mutant FUS models, including *Drosophila* larvae (Baldwin *et al.* 2016), isolated squid axoplasm (Sama *et al*. 2017), and human motor neurons derived from induced pluripotent stem cells (iPSCs) (Guo *et al.* 2017). Why then do Fus^Δ14/+^ mice not show impaired signalling endosome transport at least until a very late disease stage? In addition to the possibilities mentioned above, there are numerous potential explanations. Firstly, distinctions may arise due to the different FUS mutations being analysed. Secondly, while *Drosophila* is an excellent model that has provided instrumental insights into neurobiology, *in vivo* transport analyses are conducted in larvae in which organs have been removed, hence there is considerable disruption to the organism, which is being analysed during development and is thus perhaps not the best model for age-related neurodegeneration. Moreover, the complex, long-range neurotrophin signalling program is not conserved in *Drosophila*, while mutant ALS transgenes are often overexpressed to above physiological levels, which can induce phenotypes even with wild-type FUS transgenes (Baldwin *et al.* 2016). *In vitro* axonal transport dynamics differ from *in vivo* trafficking (Bilsland *et al.* 2010; Gibbs *et al.* 2016), possibly due to cultured neurons lacking the complete series of necessary cellular and chemical interactions (e.g. myelination and target muscle cells in the case of motor neurons) (Sleigh *et al.* 2017), which is particularly important for ALS as there are both cell and non-cell autonomous pathomechanisms (Nijssen *et al.* 2017). In addition to variability inherent to iPSC differentiation, it remains unknown how and if motor neuron developmental stages in culture correlate with age-related degeneration *in vivo*. By imaging axonal transport of signalling endosomes in intact sciatic nerves of anaesthetised mice, we can be confident of disease stage and that we are assessing transport of motor axons in their physiological environment.

In summary, we have assessed *in vivo* retrograde axonal transport of signalling endosomes in two new mouse models of ALS that express disease-causing mutant proteins at near endogenous levels. Mutant TDP-43, but not mutant Fus, mice displayed a pre-symptomatic deficiency in transport, suggesting that reduced neurotrophin signalling may contribute to mutant TDP-43-mediated neuropathology and that general defects in axonal transport are not common to all ALS-linked genes in an *in vivo* mammalian setting.

## Acknowledgements

The authors would like to thank Robert M. Brownstone (Institute of Neurology, University College London) for sharing the ChAT-eGFP mice, and the Finkelstein Laboratory (https://finkelsteinlab.org/) for the BioRxiv template.

